# Activation of the 20S proteasome core particle prevents cell death induced by oxygen- and glucose deprivation in cultured cortical neurons

**DOI:** 10.1101/2024.05.15.594129

**Authors:** Ivan L. Salazar, Michele Curcio, Miranda Mele, Rossela Vetrone, Simone Frisari, Rui O. Costa, Margarida V. Caldeira, Darci J Trader, Carlos B. Duarte

**Affiliations:** Institute for Interdisciplinary Research, University of Coimbra, Coimbra, Portugal; CNC-Center for Neuroscience and Cell Biology, University of Coimbra, Coimbra, Portugal; Multidisciplinary Institute of Ageing-MIA Portugal, University of Coimbra, Coimbra; University of California Irvine, Department of Pharmaceutical Sciences, Irvine, United States; Departament of Life Sciences, University of Coimbra, Coimbra, Portugal

## Abstract

Neuronal damage in brain ischemia is characterized by a disassembly of the proteasome and a decrease in its proteolytic activity. However, to what extent these alterations are coupled to neuronal death is controversial since proteasome inhibitors were shown to provide protection in different models of stroke in rodents. This question was addressed in the present work using cultured rat cerebrocortical neurons subjected to transient oxygen- and glucose-deprivation (OGD) as a model for in vitro ischemia. Under the latter conditions there was a time-dependent loss in the proteasome activity, determined by cleavage of the Suc-LLVY-AMC fluorogenic substrate, and the disassembly of the proteasome, as assessed by native-polyacrylamide gel electrophoresis followed by western blot against Psma2 and Rpt6, which are components of the catalytic core and regulatory particle, respectively. Immunocytochemistry experiments against the two proteins also showed differential effects on their dendritic distribution. OGD also downregulated the protein levels of Rpt3 and Rpt10, two components of the regulatory particle, by a mechanism dependent on the activity of NMDA receptors and mediated by calpains. Activation of the proteasome activity, using an inhibitor of USP14, a deubiquitinase enzyme, inhibited OGD-induced cell death, and decreased calpain activity as determined by analysis of spectrin cleavage. Similar results were obtained in the presence of two oleic amide derivatives (B12 and D3) which directly activate the 20S proteasome. Together, these results show that proteasome activation prevents neuronal death in cortical neurons subjected to in vitro ischemia, indicating that inhibition of the proteasome is a mediator of neuronal death in brain ischemia.

## Introduction

Cerebral ischemia is one of the most common causes of death and disability worldwide in developed countries (Daniele et al., 2021). The lack of oxygen and nutrients in the affected brain in stroke, the most common clinical manifestation of cerebral ischemia, leads to undesired neuronal loss through apoptosis and necrosis (Daniele et al., 2021). Accumulation of excitatory amino acids in the extracellular space, causing a toxic overactivation of extrasynaptic NMDA (N-methyl-D-aspartate) receptors (NMDAR) (excitotoxicity) (Olney, 1969; Parsons and Raymond, 2014) is a well-stablished feature in this disease. Consequently, there is a massive influx of Ca^2+^ in the postsynaptic neuron (Neves et al., 2023), which results in the overactivation of the Ca^2+^- dependent proteases calpains (Curcio et al., 2016). Transient oxygen and glucose deprivation (OGD) is a well-known protocol to model ischemia-reperfusion in cultured neurons and recapitulates many of the features that characterize neuronal degeneration in the ischemic stroke (e.g.(Mele et al., 2016)).

Conflicting results have been reported regarding the role of the Ubiquitin-Proteasome System (UPS) in neuronal damage in brain ischemia (Caldeira et al., 2014). Protein ubiquitination occurs through a three-step enzymatic cascade catalyzed by E1, E2 and E3 enzymes that target substrate proteins for degradation (Cruz Walma et al., 2022). Eight different types of polyubiquitin chain linkages can be formed, and according to this, there can be different outcomes on the ubiquitin-modified protein. For instance, it is widely accepted the K48 polyubiquitin chains are associated with degradative functions in the 26S proteasome, a 2.5 MDa multisubunit complex responsible for the controlled ATP-dependent degradation of polyubiquitinated proteins (Caldeira et al., 2014). The proteasome is composed of a catalytic 20S core particle (CP) associated with one (26S proteasome) or two (30S proteasome) 19S regulatory particles (RP), that are responsible for detecting, deubiquitinating and unfolding ubiquitinated proteins (Rousseau and Bertolotti, 2018; Sun et al., 2023).

UPS impairment has been shown to be transversal to several neurodegenerative disorders and brain ischemia (Caldeira et al., 2014; Hochrainer, 2018; Le Guerroue and Youle, 2021). The two-vessel occlusion model (15 min) induces a persistent decrease in the chymotrypsin-like activity of the proteasome when analysed up to 72 h after the insult, mirrored by an increase in polyubiquitin conjugates (Ge et al., 2007). OGD for 1.5 h also decreases the chymotrypsin-like activity of the proteasome when evaluated 4 h after the insult in cultured hippocampal neurons, and similar results were obtained when these neurons were challenged with glutamate (Caldeira et al., 2013). In addition to the decrease in proteasome activity, the 26S proteasome was found to be disassembled into its major constituents, the 19S and the 20S particles, in models of transient forebrain ischemia and global ischemia (Asai et al., 2002; Ge et al., 2007), and in cultured neurons subjected to excitotoxic glutamate stimulation (Caldeira et al., 2013). In vitro, it was also observed an accumulation of polyubiquitinated proteins in cerebrocortical neurons exposed to OGD for 2 h followed by different periods of post-incubation period under normoxic conditions (Chen et al., 2010). The identity of these proteins was resolved and were found to be involved in important neuronal functions and signalling pathways (Iwabuchi et al., 2014). More recent studies of phospho-proteomics revealed altered PSD-associated phosphorylation patterns that are regulated by ubiquitin during ischemic stroke (Dhawka et al., 2024). However, there are conflicting data linking proteasome activation and/or inhibition to neuroprotection in the context of brain ischemia. CVT-634, the first proteasome inhibitor to be tested in a rat model of focal brain ischemia, reduced infarction without affecting regional cerebral blood flow (Buchan et al., 2000). Similarly, the effect of the proteasome inhibitor MLN519 was tested using transient middle cerebral artery occlusion (MCAo) (Williams et al., 2004; Williams et al., 2005), and the cardioembolic stroke model (Zhang et al., 2001). In the former model, a therapeutic window of 6-10 h after ischemia/reperfusion brain injury was observed. The proteasome inhibitor Bortezomib was also shown to reduce infarct volume and neurological functional deficit when administrated within 4 h after stroke onset (Zhang et al., 2006). Systemic proteasome inhibition with BSc2118 also induced sustained post-stroke neurological recovery and neuroprotection, when delivered up to 9 h after stroke (Doeppner et al., 2016; Doeppner et al., 2012). In these studies, the effect of proteasome inhibitors was mainly attributed to stabilization of the blood-brain barrier integrity (Caldeira et al., 2014). A dichotomic effect of proteasome inhibition/activation is supported by studies showing that IU-1, an USP14 inhibitor and therefore a UPS enhancer (Lee et al., 2016), reduced brain infarct volume in a mouse model of transient focal cerebral ischemia at day 4 after the insult (Doeppner et al., 2013). The USP14 inhibitor IU-1 promotes the degradation of the transcription factor REST (Doeppner et al., 2013). Similar work also showed that IU-1 improves survival and functional recovery, reduces infarct volume, and restores ubiquitin homeostasis in the transient MCAo model of focal brain ischemia (Min et al., 2017). Yet, the molecular mechanisms associated with this improvement were not addressed. In addition, to what extent the alterations in neuronal proteasome structure and function observed in brain ischemia contribute to neuronal demise remain to be investigated.

In this study, we sought to investigate the molecular mechanisms contributing to the alterations in proteasome structure and functions in neurons during brain ischemia, and how alterations in the proteasome activity contribute to neuronal damage. Studies were performed in cultured rat cerebrocortical neurons transiently exposed to OGD and showed a role for calpains in the selective cleavage of proteasome subunits. The use of cultured neurons allowed to avoid interfering with the role of the proteasome in the stability of the blood brain barrier (see above). Activation of the 20S proteasome in cultured neurons subjected to OGD significantly decreased calpain activation and reduced neuronal death, suggesting a crosstalk between the two proteolytic systems.

## Materials and Methods

### Cerebrocortical cultures (high-density and low-density cultures)

Primary cultures of rat cortical neurons were prepared from the cortices of E17-E18 Wistar rat embryos, as previously described (Salazar et al., 2017). Briefly, cortices were washed with ice-cold HBSS three and five times, prior and after trypsin (0.06 %, 10 min at 37°C) treatment, respectively. Cells were mechanically dissociated with a 5 ml glass pipette, no more than 10-15 times with HBSS. After counting, the cells were plated with Neuronal Plating Medium (MEM supplemented [Sigma-Aldrich] with 10 % horse serum [ThermoFisher], 0.6 % glucose and 1 mM pyruvic acid) for 2-3 h in 6- or 24-well plates (92.8x10^3^ cells/cm^2^) coated with poly-D-lysine (0.1 mg/mL). After this period, the plating medium was removed and replaced by Neurobasal medium supplemented with SM1 supplement (StemCell Technologies; 1:50 dilution), 0.5 mM glutamine and 0.12 mg/mL gentamycin. For imaging purposes, low-density cortical cells were plated at a final density of 1.0x10^4^ cells/cm^2^ on poly-D-lysine-coated coverslips, in 60 mm culture dishes, in neuronal plating medium (Mele et al., 2021). After 2-3 h, coverslips were flipped over an astroglial feeder layer in supplemented Neurobasal medium, and upon 2-3 days in culture the division of glial cells was halted by addition of 10 µM 5-FdU-NOAC (5-FDU) to the culture medium. Cultures were fed twice a week and maintained in Neurobasal medium supplemented with SM1 supplement, and kept in a humidified incubator of 95% air and 5 % CO_2_, at 37°C, for 14-15 days.

### Oxygen-Glucose Deprivation (OGD) Assays

Cultured cerebrocortical neurons were incubated for 45 min or 1h30min (see figure captions) in a solution containing 10 mM Hepes, 116 mM NaCl, 5.4 mM KCl, 0.8 mM MgSO_4_, 1 mM NaH_2_PO_4_, 25 mM NaHCO_3_, 1.8 mM CaCl_2_, pH 7.3, and supplemented with 25 mM glucose (Sham) or with 25 mM sucrose (OGD) at 37°C, as previously described (Caldeira et al., 2014; Mele et al., 2016).

In the Sham condition, cells were maintained in an incubator with 5 % CO_2_/95 % air, whereas the latter group was incubated in an oxygen deprived chamber (5 % CO_2_, 7.5 % H_2_, 87.5 % N_2_; OGD condition) (Thermo Forma Anaerobic System Model 1029). Cells were further incubated in culture conditioned medium for 4 h (for preparation of protein extracts), or for the period of time indicated in figure captions (for measurement of the proteasome activity and cell death experiments). When appropriate, 50 µM MDL 28170 (Calbiochem) or 100 µM APV (Enzo Lifesciences) were added to the culture medium 30 min before OGD and were present throughout the experimental procedure. The DUB USP14 inhibitor IU1 (20 µM; Focus Biomolecules) was added to the culture medium 30 min before OGD, or immediately after, and was present throughout the experiment. The 20S oleic derivatives B12 and D3 were synthesized as previously reported (Halder et al., 2022) and were added to the culture medium 2 h before OGD (10 µM), and were present throughout the experiment.

### Immunocytochemistry

Cultured cerebrocortical neurons (low-density cultures) were fixed in 4 % sucrose/4 % paraformaldehyde (in PBS) for 15 min at room temperature and permeabilized with 0.3 % Triton X-100 in PBS. Neurons were then incubated with 10 % BSA in PBS, for 30 min at 37°C, to block non-specific staining, and incubated overnight at 4°C with the primary antibodies diluted in 3 % BSA in PBS. The following primary antibodies and dilutions were used: anti-Psma2 (1:300 dilution [rabbit]; Cell Signaling #2455), anti-Rpt6 (1:400 dilution [mouse]; Enzo #BML-PW9265), anti- MAP2 (1:10,000 dilution [chicken], Abcam #ab5392), anti-PSD95 (1:200 dilution [rabbit]; Cell Signaling # 1673450S) and anti-PSD95 (1:200 [mouse]; Thermo Scientific #MA1-045). The cells were washed 6 times with PBS for 2 min and incubated with Alexa Fluor 568- (1:500 dilution, Invitrogen), Alexa Fluor 488- (1:500 dilution; Invitrogen) and AMCA- (1:200 dilution; Jackson ImmunoResearch) conjugated secondary antibodies, for 45 min at 37°C. After washing the cells 6 times with PBS for 2 min, the coverslips were mounted with a fluorescence mounting medium (DAKO).

### Flurorescence microscopy and quantitative fluorescence analysis

Imaging was performed in an Axio Observer Z1 fluorescence microscope, coupled to an Axiocam HRm digital camera, using a Plan-Apochromat 63x/1.4 oil objective. Images were quantified using the ImageJ image analysis software. For quantitation, independent sets of cells were cultured and stained simultaneously, and imaged using identical settings. The immunoreactivity signals were analysed after setting the thresholds, and the recognizable clusters under those conditions were included in the analysis. The number, area and the integrated intensity of Psma2 or Rpt6 particles in dendrites were determined and represented per dendritic area. For colocalization analysis, regions around thresholded puncta were overlaid as a mask in the PSD95 (or MAP2) channel, and the integrated intensity, area and number of colocalized particles determined. Fluorescence imaging was performed at the MICC Imaging facility of CNC-UC, partially supported by PPBI – Portuguese Platform of BioImaging (PPBI-POCI-01-0145-FEDER- 022122).

### Nuclear morphology staining

Cerebrocortical neurons were cultured for 14 days on poly-D-lysine-coated glass coverslips, at a density of 1.0x10^4^ cells/cm^2^. After the appropriate stimulus (see figure captions), cells were fixed in 4 % sucrose/ 4 % paraformaldehyde (in PBS), for 15 min at room temperature, washed twice with ice-cold PBS and the nuclei were then stained with Hoechst 33342 (1 µg/mL), for 10 min, protected from the light and at room temperature. After this, cells were washed twice with ice-cold PBS and the coverslips were mounted with a fluorescent mounting medium (DAKO). Images were captured using a Zeiss Axiovert 200 fluorescent microscope coupled to an Axiocam camera. Three independent coverslips were prepared for each experimental condition, and at least 200 cells were counted in two biological replicates for each experimental condition.

### Preparation of extracts and quantification of proteasome activity

Cultured cerebrocortical neurons were washed twice with ice-cold phosphate-buffered saline (PBS). The cells were then lysed in 1 mM EDTA, 10 mM Tris-HCl pH 7.5, 20 % Glycerol, 4 mM dithyothreitol (DTT) and 2 mM ATP (100 µl/well). Whole cell extracts were centrifuged at 16,100 x g for 10 min at 4°C, total protein content in the supernatants was quantified using the Bradford method, and the concentration of the samples was equalized with lysis buffer.

The peptidase activity of the proteasome was assayed by monitoring the production of 7-amino- 4-methylcoumarin (AMC) from a fluorogenic peptide: Suc-LLVY-AMC (for chymotrypsin-like activity; Peptide Institute, Inc). Samples (5-10 µg of protein) were incubated with the fluorogenic substrate, 50 µM Suc-LLVY-AMC, in 50 mM Tris-HCl (pH 8.0) and 0.5 mM EDTA buffer, in a final volume of 100 µl. The release of fluorescent AMC was measured at 37°C using a SPECTRAmax Gemini EM (Molecular Devices) microplate reader, at an excitation wavelength of 360 nm and an emission wavelength of 460 nm, for 60 min at 5 min intervals. All experiments were performed in the presence of 2 mM ATP. Specific activity was determined by subtracting the

activity measured in the presence of 10 µM MG-132 (Calbiochem), a reversible proteasome inhibitor.

### Native Gel Electrophoresis

Cultured cerebrocortical neurons were washed twice with ice-cold PBS and the cells were then lysed in 100 µl of 50 mM Tris-HCl pH 7.5, 10 % Glycerol, 5 mM MgCl_2_, 2 mM DTT and 2 mM ATP. The extracts were then centrifuged at 16,100 x g for 10 min at 4°C, and the protein content in the supernatants was quantified using the Bradford method. The protein concentration in the samples was equalized with lysis buffer before separation in 4 % polyacrylamide native gels, under non-denaturing conditions, at 90 V for 8 h (4°C) as described elsewhere (Elsasser et al., 2005; Caldeira et al., 2013). The proteins in the gel were then electrotransferred to polyvinylidene (PVDF) membranes and immunoblotted using an antibody against Psma2 (1:1000 dilution [rabbit]; Cell Signaling #2455) or Rpt6 (1:1000 dilution [mouse]; Enzo #BML-PW9265).

### Western Blotting

Cultured cerebrocortical neurons were washed with ice-cold PBS buffer and then lysed with RIPA buffer (150 mM NaCl, 50 mM Tris-HCl, pH 7.4, 5 mM EGTA, 1 % Triton, 0.5 % DOC and 0.1 % SDS at a final pH 7.5) supplemented with 50 mM NaF, 1.5 mM sodium orthovanadate and the cocktail of protease inhibitors (CLAP [1 µg/ml]: chymostatin, leupeptin, antipain, pepstatin; Sigma)]. After centrifugation at 16,100 x g for 10 min, protein in the supernatants was quantified using the bicinchoninic acid (BCA) assay, and the samples were diluted with a 2x concentrated denaturating buffer (125 mM Tris, pH 6.8, 100 mM glycine, 4 % SDS, 200 mM DTT, 40 % glycerol, 3 mM sodium orthovanadate, and 0.01 % bromophenol blue).

Protein samples were separated by sodium dodecyl sulfate polyacrylamide gel electrophoresis (SDS-PAGE), in 6.5-11 % polyacrylamide gels, transferred to PVDF membranes (Millipore), blocked for 45 min, with 5 % non-fat milk in Tris-buffered saline supplemented with 0.1% Tween 20 (TBS-T) and immunoblotted using commercial antibodies. Blots were incubated with primary antibodies (overnight at 4°C), washed and exposed to alkaline phosphatase-conjugated secondary antibodies (1:20,000 dilution; Jackson Immunoresearch; 1 h at room temperature in TBS-T). The following primary antibodies were used: Anti-Psma2 (1:1000 dilution [rabbit]; Cell Signaling #2455), Anti-Rpt6 (1:1000 dilution [mouse]; Enzo #BML-PW9265), Anti-Rpn10 (1:1000 dilution [mouse]; Enzo # BML-PW9250), Anti-Spectrin (1:1000 dilution [mouse]; Millipore # MAB1622), Anti-Rpt3 (1:100 dilution [mouse]; Santa Cruz Biotechnology #sc-166115), Anti-Rpn6 (1:1000 dilution [rabbit]; Cell Signaling # 14303). Alkaline phosphatase activity was visualized by ECF on the Storm 860 Gel and Blot Imaging System (GE Healthcare) or the ChemiDoc Touch Imaging System (BioRad).

To reprobe membranes with additional primary antibodies, namely those used against β- tubulin, used as experimental loading control, ECF was removed by washing the membranes with TBS-T for 30-40 min. After this washing step, the membranes were stripped for 5 min with 0.2 M NaOH, and washed again abundantly with water for 30 min. Membranes were again blocked and incubated with the anti-β-tubulin antibody (1:600 000). Secondary antibodies were incubated for 1 h at room temperature as previously mentioned.

### Lactate dehydrogenase activity

After the OGD insult, the cells were further incubated in culture conditioned medium for the period of time mentioned in the figure captions. The LDH leakage to the extracellular medium was evaluated by a colorimetric assay, using the CytoTox 96® Non-Radioactive assay kit (Promega), according to the manufactureŕs instructions. Briefly, the extracellular medium was removed and diluted with an equal volume of H_2_O to a final volume of 100 μl, and 50 μl of substrate mix was added to the diluted sample, at room temperature, protected from the light. Incubation with the substrate was performed for 15-30 min and the reaction was stopped with 50 μl of stop solution. The activity of LDH was measured using a SPECTRAmax Gemini EM (Molecular Devices) microplate reader, at an wavelength of 490 nm. The percentage of LDH released to the medium was determined as the ratio between LDH activity in the extracellular medium and total LDH activity (100 % cell death) obtained by cell lysis.

### Statistical Analysis

Statistical analysis was performed using one-way ANOVA analysis of variance followed by the Dunnett’s or Bonferroni test, or using the two-tailed Student’s *t* test, as indicated in the figure captions.

## Results

### OGD decreases proteasome activity and induces its disassembly

OGD is a well-characterized approach to model global brain ischemia-reperfusion in cultured neurons (Caldeira et al., 2013; Costa et al., 2016; Curcio et al., 2015; D’Orsi et al., 2015; Mele et al., 2016). To characterize the effect of OGD on proteasome activity, cerebrocortical neurons were transiently subjected to OGD for 1.5 h and the chymotrypsin-like activity of the proteasome was evaluated after further incubation of the cells for different time periods in culture conditioned medium in normoxic conditions. Exposure of cerebrocortical neurons to OGD for 1.5 h rapidly decreased the activity of the proteasome, to 47 - 62 % of the control (Figure 1A; *0.05<p<0.0001****). Importantly, under these conditions there was an increase in cell death from 14 to 27 %, as determined by DNA staining with Hoechst 33342 at 10 h after the insult (Figure 1B; **p<0.01). Cell death was characterized by chromatin condensation and a decrease in the size of nuclei, which is characteristic of apoptotic-like death and is in accordance with previous findings under the same conditions (e.g. (Mele et al., 2016)).

**Figure 1.**
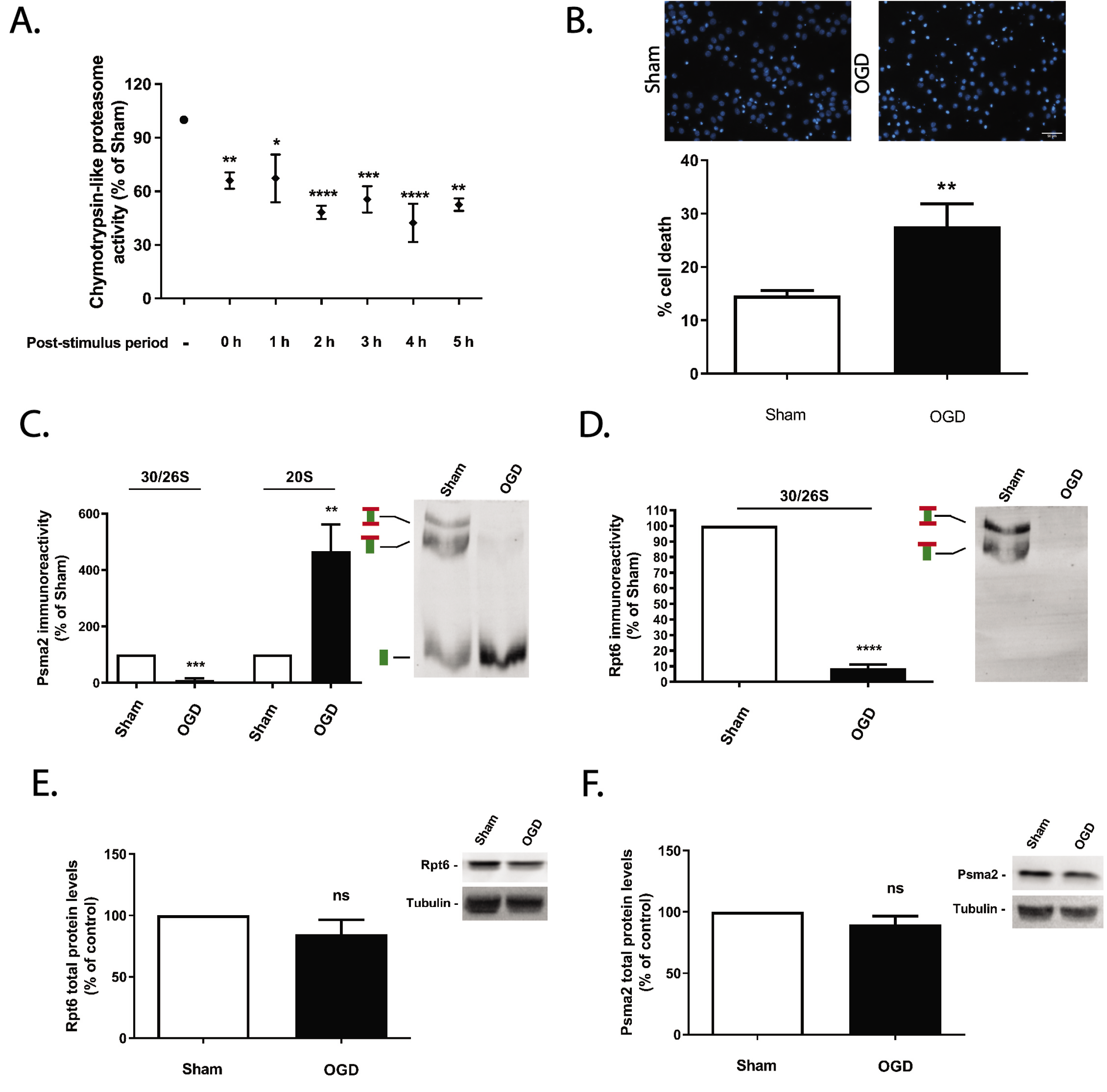
Effect of OGD on the activity of the proteasome and its assembly in cerebrocortical neurons. Cerebrocortical neurons were subjected to OGD for 1.5 h and were further incubated in culture conditioned medium for the indicated periods of time. (A) The chymotrypsin-like activity of the proteasome was measured with the fluorogenic substrate 25 µM Suc-LLVY-AMC and cell death was evaluated by nuclear morphology analysis (B), staining the nucleus with Hoechst 33342, and presented in the quantification graph as the percentage of cell death. The scale bar represents 10 µm. The control proteasome activity, determined in neurons not exposed to OGD, was set to 100%. The abundance of Psma2 (C) and Rpt6 (D) was determined under non-denaturating conditions after immunoblotting the native gel (Native-PAGE) with antibodies against Psma2, to probe the 26S/30S and the 20S proteasomes, and against Rpt6, to identify 26S proteasomes. Total abundance of Psma2 (E) and Rpt6 (F) was also evaluated under denaturating conditions (SDS-PAGE) as control experiments, and β-tubulin was used as loading control. Control (Sham) protein levels of Pmsa2 or Rpt6 subunits were set to 100 %. The results are the average ± SEM of 3-4 independent experiments. Statistical analysis was performed by the Student’s *t* test (ns, p>0.5; *p˂ 0.5; **p˂ 0.01 ***p˂ 0.001; ****p˂ 0.0001).

In additional experiments we compared the alterations in the assembly of the proteasome in cultured cortical neurons subjected to OGD for 1.5 h, and further incubated in culture conditioned medium for 4 h; these are the conditions that showed the most robust downregulation in chymotrypsin-like activity of the proteasome. We observed a reduction in the population of fully-assembled proteasomes (26S and 30S, for single- and double-capped proteasomes, respectively) to about 10 %, as shown by a decrease in the catalytic core particle Psma2 immunoreactivity in the high molecular weight fraction (10 %) (Figure 1C, left panel; ***p<0.001), as well as in the regulatory particle protein Rpt6 (8 %) (Figure 1D ****p<0.0001). Similarly, the 20S proteasomes were upregulated to more than 4-fold, as shown by an increase in the Psma2 immunoreactivity in the low molecular weight fraction (Figure 1C right panel; **p<0.01). Control western blot experiments to evaluate total levels of Psma2 and Rpt6 proteins, showed no changes in total abundance of both proteins, further indicating a proteasome disassembly under these conditions (Figure 1E and F; ns p>0.05; Figure 1. Supplementary information).

Together, these data show a proteasomal dysfunction following OGD, associated to the disassembly of the complex.

### Transient OGD alters proteasome subcellular localization during reperfusion

The UPS is responsible for the turnover of many cytosolic proteins, and under resting conditions their components show a widespread distribution in neurons, including nuclei, cytoplasm, dendrites, axons and synaptic buttons (Mengual et al., 1996). Little is known about the proteasome dynamics after brain ischemia, such as the alterations in the subcellular localization of the two components of the 26S proteasome. Therefore, we evaluated the effect of OGD on the distribution of the proteasome in low density cultures of cerebrocortical neurons during different reperfusion periods. We first adapted the OGD protocol to low density cultures (Banker cultures; 1.0x10^4^ cells/cm^2^) (Kaech and Banker, 2006) since high density cultures do not allow assessing accurately the subcellular distribution of proteins using immunocytochemistry. Given the putative protective effect provided by glial cells when present during the period of OGD, we tested the ischemic insult on neurons in the absence of the glial cell feeder layer. After OGD, cortical neurons were further incubated in culture conditioned medium and in the presence of glial cells. A significant increase in cell death was observed after ischemic insults of 45 min (∼30 %) when analysed 4 h (*p<0.05) and 8 h (**p<0.01) after the OGD period (Figure 2). Therefore, to facilitate subsequent co-localization experiments, we employed a 45-min OGD period followed by several post-incubation periods. The briefer OGD duration was chosen due to its association with minimal cell death and no change in the pattern of staining for the dendritic marker (see below).

**Figure 2.**
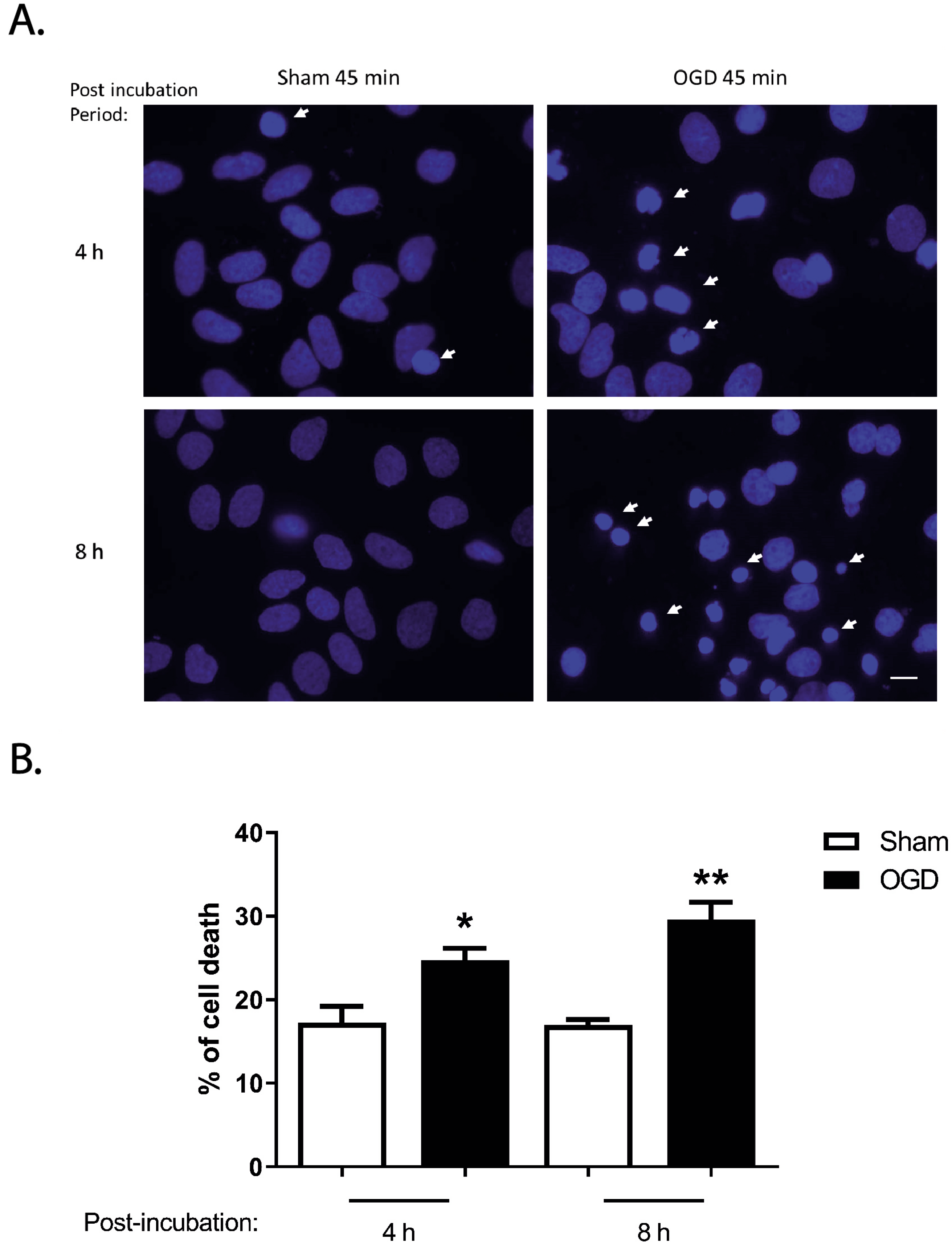
Effect of OGD on neuronal survival in low-density cultures. Low-density cultured cerebrocortical neurons were subjected or not to OGD for 45 min, and further incubated in culture conditioned medium for 4 and 8 h. Cell death was evaluated by nuclear morphology analysis (A), staining the nucleus with Hoechst 33342, and presented in the quantification graph as the percentage of cell death (B). Arrows point to dead cells. The scale bar represents 10 µm. The results are the average ± SEM of 4-5 independent experiments. Statistical analysis was performed by one-way ANOVA, followed by Bonferronís multiple comparison test, comparing selected pairs of columns (*p˂ 0.05; **p˂ 0.01).

Despite the clear heterogeneity in their subunit composition, it is possible to label the two proteasomal particles, the 20S and the 19S, by means of antibodies recognizing Psma2 and Rpt6 proteins, respectively (Sun et al., 2023). Both proteins are found in the soma and dendrites of cultured neurons as evaluated by the colocalization with the somato-dendritic marker MAP2 (Figures 3 and 4). The results of Figure 3 show a significant increase of the area (Figure 3A2, 1.42- fold; ****p<0.0001) and intensity (Figure 3A3, 1.62-fold; ****p<0.0001) of Psma2 puncta in dendrites after 45 min of OGD followed by 2 h of post-incubation. These effects were transient since 4 h after OGD these differences were no longer observed. At a later time-point, i.e. 6 h after OGD, there was a secondary increase in the puncta number (Figure 3A1, 1.25; *p<0.05) and intensity (Figure 3A3, 1.99-fold; **p<0.01), without further changes in the area of the puncta. Further analysis of the OGD-induced changes in the Psma2 distribution along dendrites showed a significant, although transient, increase in the number (Figure 3A4, 1.41-fold; ***p<0.001) and area (Figure 3A5, 1.52-fold; **p<0.01) of puncta that colocalized with the post- synaptic density marker PSD95, when analysed 2 h after the ischemic insult. In addition, the percentage of total Psma2 immunoreactivity that colocalized with PSD95 was also increased at 2 h after OGD, but no similar effects were observed when the Psma2 distribution was analysed at later time points after the ischemic insult (Figure 3A6, 1.24-fold; **p<0.01).

**Figure 3.**
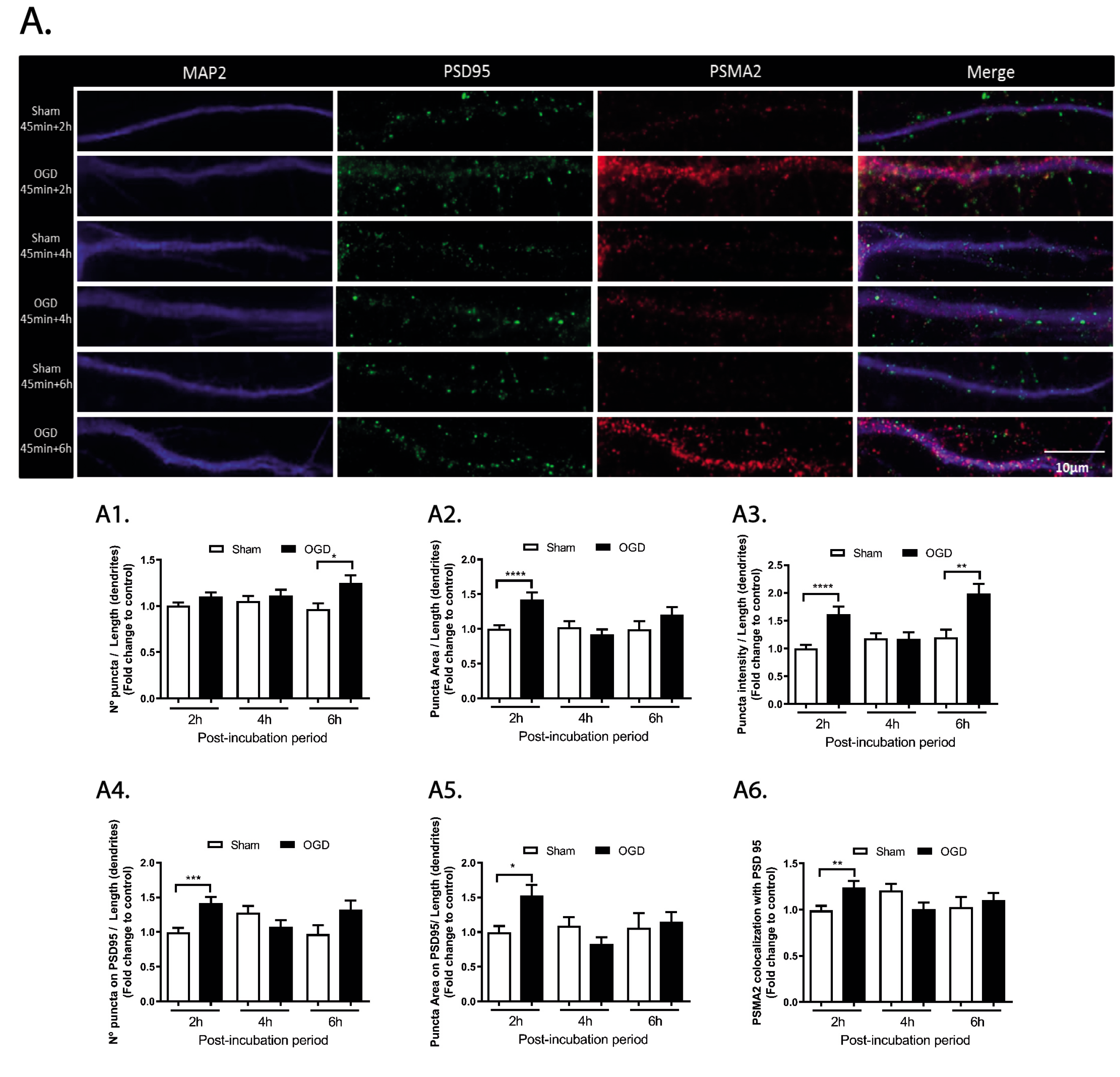
Alterations in the dendritic distribution of PSMA2 in cerebrocortical neurons subjected to OGD. PSMA2 protein expression was evaluated by immunocytochemistry in neurons subjected to 45 min of OGD and further incubated in culture conditioned medium for 2, 4 or 6 h. The number (A1), area (A2), and intensity (A3) of PSMA2 clusters (puncta) were analyzed in the dendritic compartment through colocalization with MAP2, and normalized for its length. The number (A4) and area (A5) of PSMA2 puncta colocalizing with PSD95, as well as the percentage of PSMA2 puncta colocalized with PSD95 (normalized for the mean of the control) (A6), were also calculated. Imaging was performed in an Axio Observer Z1 fluorescence microscope, coupled to an Axiocam HRm digital camera, using a Plan-Apochromat 63x/1.4 oil objective. The scale bar represents 10 µm. Results are mean ± SEM of at least 3 independent experiments (30 cells). Statistical analysis was performed by one-way ANOVA, followed by Bonferroni test. *p<0.05; **p<0.01; ***p<0.001; ****p<0.0001 - significantly different when compared to control conditions.

**Figure 4.**
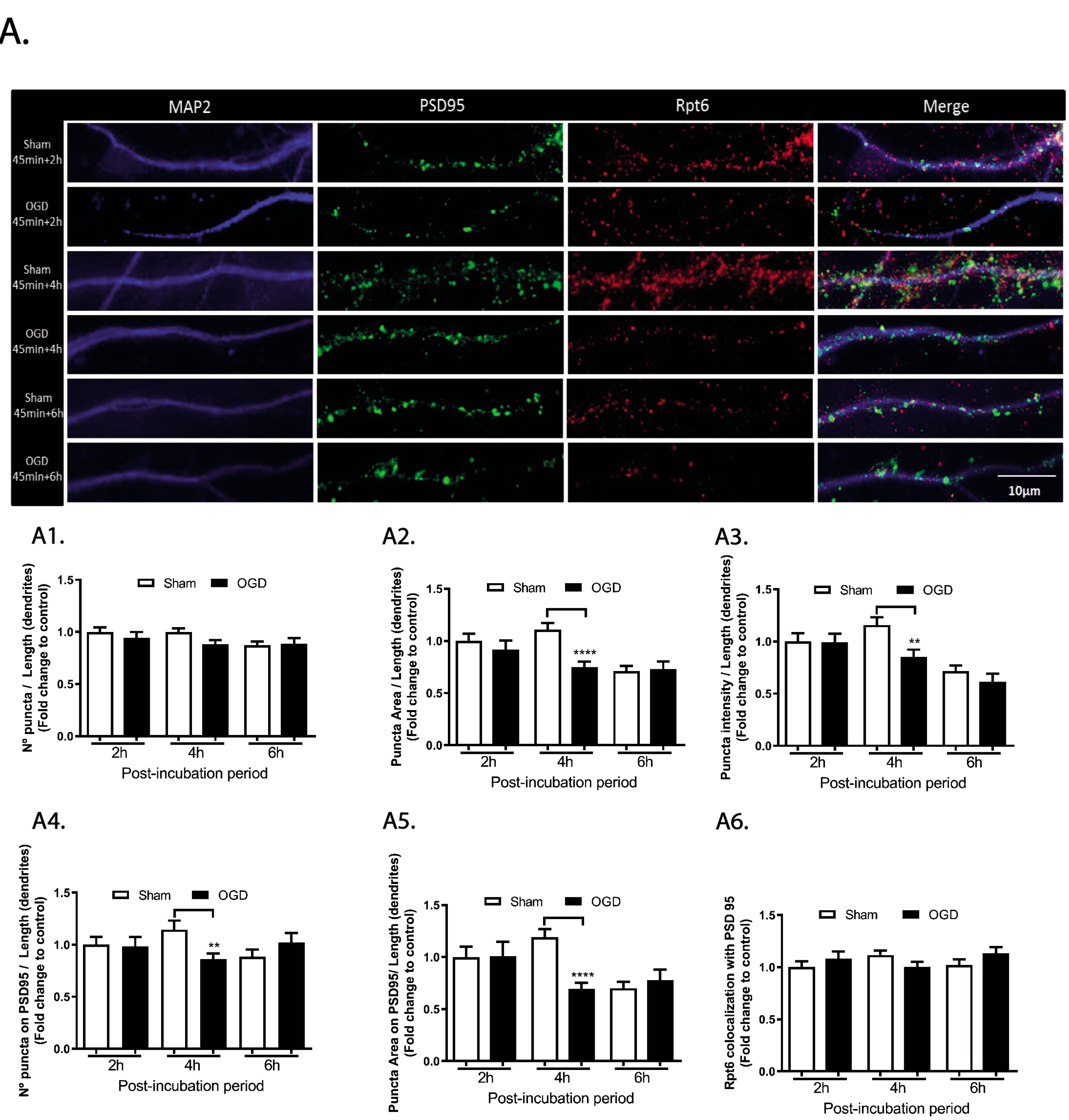
Alterations in the dendritic distribution of Rpt6 in cerebrocortical neurons subjected to OGD. Rpt6 protein expression was evaluated by immunocytochemistry in neurons subjected to 45 min of OGD and further incubated in culture conditioned medium for 2, 4 or 6 h. The number (A1), area (A2), and intensity (A3) of Rpt6 clusters (puncta) were analyzed in the dendritic compartment through colocalization with MAP2, and normalized for its length. The number (A4) and area (A5) of Rpt6 puncta colocalizing with PSD95, as well as the percentage of Rpt6 puncta colocalized with PSD95 (normalized for the mean of the control) (A6), were also calculated. Imaging was performed in an Axio Observer Z1 fluorescence microscope, coupled to an Axiocam HRm digital camera, using a Plan-Apochromat 63x/1.4 oil objective. The scale bar represents 10 µm. Results are means ± SEM of at least 3 independent experiments (30 cells). Statistical analysis was performed by one-way ANOVA, followed by Bonferroni test. **p<0.01; ****p<0.0001 - significantly different when compared to control conditions.

In contrast with the observations for the Psma2 protein, OGD did not affect the number (Figure 4A1), area (Figure 4A2) and intensity (Figure 4A3) of Rpt6 puncta along dendrites when evaluated 2 h and 6 h after the insult (ns p>0.05). Furthermore, no changes were detected in the number and area of Rpt6 puncta that colocalized with PSD95 (Figure 4A4-A5), as well as in the total protein fraction that colocalized with PSD95 (Figure 4A6). Alterations in the subcellular distribution of Rpt6 were only observed at 4 h after the insult, with a decrease in the area (Figure 4A2, 0.74-fold; ****p<0.0001) and intensity of puncta (Figure 4A3 0.85-fold; **<0.01), as well as in the puncta number (Figure 4A4, 0.86-fold; **p<0.01) and area (Figure 4A5, 0.69-fold; ****p<0.0001) that colocalized with PSD95.

Together, these results show differential effects of OGD in the localization Psma2 and Rpt6, which were used to label the 20S and 19S proteasome, respectively. The distribution of Psma2 was more affected at 2 h after the insult, while the localization of Rpt6 only showed changes at 4 h after OGD.

### OGD induces calpain activation: a possible link leading to proteasome dysfunction

The abundance of the proteasome can be regulated by several ways, in cells destined to die. For instance, apoptotic Jurkat T cells showed a cleavage of several proteasome subunits by caspase- 3, such as Rpt5, Rpn2, Rpn1, Rpn10 subunits (Adrain et al., 2004; Sun et al., 2004), while α2, α4, ß4 and Rpt1 were truncated by caspase-7 (Jang et al., 2007). Furthermore, Rpn10 can be cleaved by calpains in the presence of mitochondrial toxins in cultured cortical neurons (Huang et al., 2013). Therefore, we hypothesized that calpains, a group of Ca^2+^-dependent cysteine proteases, could play a role in the inactivation of the proteasome in ischemic conditions, by limited proteolysis of specific proteasomal subunits. Indeed, the [Ca^2+^]_i_ overload followed by activation of calpains is a hallmark of ischemia/reperfusion, both in in vivo and in vitro models, as is also characteristic of NMDAR-dependent excitotoxicity (Curcio et al., 2015; Neumar et al., 2001).

Cerebrocortical neurons subjected to OGD for 1.5 h exhibited a 4-fold increase in the calpain-mediated cleavage of spectrin (a well-known hallmark of calpain activation), giving rise to the spectrin breakdown products (SBDP) 150/145, when compared to the control (Figure 5A;^++++^p<0.0001; Figure 5. Supplementary information). The abundance of SBDP150/145 was reduced to basal levels in the presence of the calpain inhibitor MDL28170 (Figure 5A; ns p>0.05, when compared with the control), or NMDAR antagonist APV (Figure 5A; ns p>0.05). Both inhibitors completely abrogated the increase of SBDP150/145 observed in cortical neurons subjected to OGD (Figure 5A; ****p<0.0001). Furthermore, in the absence of the inhibitors there was no apparent formation of the SBDP120, further suggesting that caspase-3 is not a key mediator of cellular responses under these experimental conditions (Wang, 2000).

**Figure 5.**
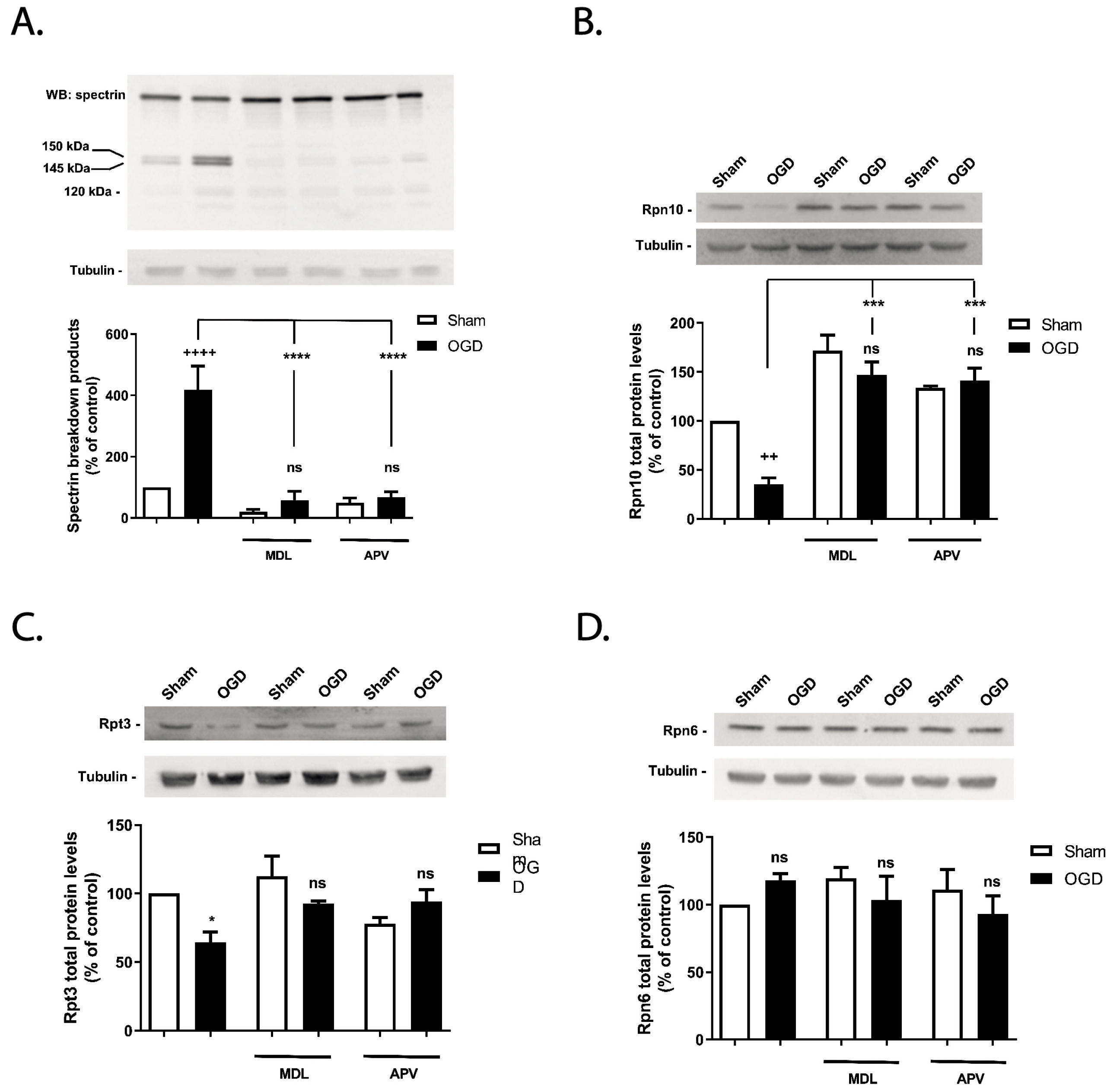
Effect of OGD on calpain activation and on Rpn6, Rpn10 and Rpt3 total protein levels in cultured cerebrocortical neurons. Cultured cerebrocortical neurons were subjected to OGD for 1.5 h and were further incubated in culture conditioned medium for 4 h. When appropriate, cerebrocortical neurons were incubated with 100 μM APV or 50 μM MDL28170. The inhibitors were added 30 min prior and during OGD, and were also present during the post-incubation period. (A) Spectrin full-length (280 kDa) and SBDPs (150 and 145 kDa) were analyzed by western blot using an antibody recognizing both the N- and C-terminal regions of the protein. The total abundance of (B) Rpn10, (C) Rpt3 and (D) Rpn6 proteins levels were also evaluated by immunoblot. β-tubulin was used as loading control. Control (Sham without MDL or APV) protein levels of spectrin were set to 100 %. The results are the average ± SEM of 3 independent experiments. Statistical analysis was performed by one-way ANOVA, followed by the Bonferronís multiple comparison test, comparing all the conditions with the respective control (ns, p>0.5; ^++++^p˂ 0.0001; *p˂ 0.5; ***p˂ 0.001 ****p˂ 0.0001).

We further investigated the role of calpains in the cleavage of several proteasome subunits in cultured cerebrocortical neurons exposed to OGD or maintained under control conditions. OGD (1.5 h) significantly decreased (35 %) Rpn10 protein levels (a regulatory subunit) in cultured cortical neurons, when evaluated after 4 h of reoxygenation (Figure 5B; ^++^p<0.001; Figure 5. Supplementary information), but the antibody used did not detect the truncated protein (not shown). The effect of OGD was abrogated in the presence of the calpain inhibitor MDL28170 (Figure 5B; ns p>0.05), and by the NMDAR antagonist APV (Figure 5B; ns p>0.05). Similarly, we observed a decrease of about 36 % in total Rpt3 protein abundance (Figure 5C; *p<0.01). This decrease was prevented when neurons were pre-incubated with the calpain inhibitor MDL28179 (Figure 5C; ns p>0.05; Figure 5. Supplementary information), and with the NMDAR antagonist APV (Figure 5C; ns p>0.05). In contrast, we did not find any differences in Rpn6 total proteins levels in neurons subjected to OGD (Figure 5D; ns p>0.05; Figure 5. Supplementary information), showing the specificity of the effects of calpains on the cleavage of some proteasome subunits.

Altogether, the results obtained show a calpain-mediated selective cleavage of 26S proteasome subunits under conditions that mimic brain ischemic in vitro. However, due to the heterogeneity of the complex, we cannot definitively exclude the possibility of cleavage occurring in other subunits. Further studies are required to address whether and to what extent other proteasomal subunits are cleaved by calpains under excitotoxic conditions.

### Proteasome activation as a protective strategy to prevent cell death evoked by OGD in cultured cerebrocortical neurons

Proteasome downregulation as well as its disassembly is a common feature observed in models of transient global/focal ischemia, and also in in vitro models (Asai et al., 2002; Caldeira et al., 2013; Ge et al., 2007; Hochrainer et al., 2012; Hu et al., 2000). However, these observations contrast with the neuroprotective effects resulting from proteasome inhibition in several in vivo models of brain ischemia (e.g (Doeppner et al., 2016; Doeppner et al., 2012; Williams et al., 2004; Williams et al., 2003)). Whether inhibition of the proteasome contributes to neuronal death in cerebrocortical neurons exposed to OGD was assessed using two different approaches: (i) IU1 is an inhibitor of USP14, a DUB associated with the 26S proteasome, and was shown to enhance the degradation of mutant proteins associated with several neurodegenerative disorders such as Tau, Ataxin-3 and TDP-43 in cell lines (Lee et al., 2010); (ii) activators of the 20S core particle.

First, cerebrocortical neurons were subjected to OGD for 1.5 h, and the neuroprotective effect of the UPS activator IU1 (20 μM) was evaluated by analyzing the nuclear morphology 10 h after the insult. The concentration of IU1 used here is within the range of concentrations that affect the degradation of endogenous substrates without compromising cell viability (up to 100 μM) in human-derived cell lines (Lee et al., 2010; Nag and Finley, 2012). Pre-incubation of the cerebrocortical neurons with IU1 abrogated OGD-induced cell death, as determined by analysis of the nuclear morphology (Figure 6A,B; ns p>0.05). However, the protective effect of IU1 was not observed when the drug was applied immediately after the insult (post-OGD). The results obtained under the latter conditions are comparable to those obtained in vehicle-treated cells (Figure 6A,B; *p<0.05). Similar results were obtained when the release of LDH, a cytosolic enzyme, to the extracellular medium was evaluated. We observed a neuroprotective effect of IU1 when applied 30 min before the insult, and a loss of this effect when the drug was added immediately after the OGD insult (Fig 6C; ns p>0.05 for pre-OGD, and *p<0.05 for post-OGD). To investigate whether incubation with IU1 affects calpain activation following OGD, the activity of the protease was analyzed by measuring the cleavage of spectrin (Figure 6D; Figure 6. Supplementary information). Cerebrocortical neurons exposed to OGD for 1.5 h and further incubated in culture conditioned medium for 4 h showed a 1.9-fold increase of the SBDP150/145 (Figure 6D; *p<0.05), which is indicative of calpain activation (Wang, 2000). However, when 20 μM of IU1 was present there was no increase in SBDPs 150/145 kDa when the Sham and OGD conditions were compared, indicating that there was no calpain activation (Figure 6D; ns p>0.05). These results suggest that proteasome activation inhibits OGD-induced upregulation in calpain activity.

**Figure 6.**
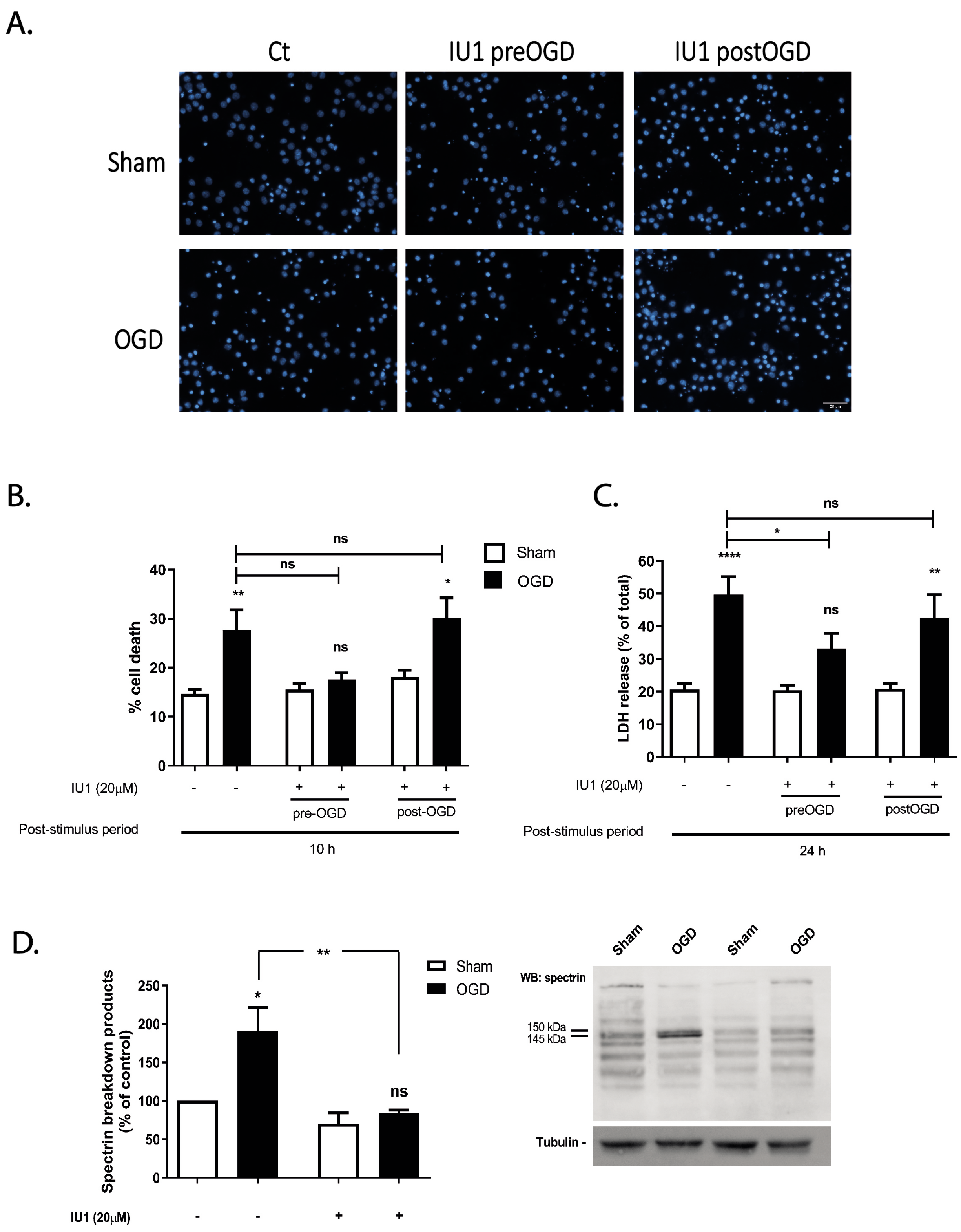
Cell death evoked by OGD is prevented by the USP14 inhibitor IU1. Cultured cerebrocortical neurons were subjected or not to OGD for 1.5 h, and further incubated in culture conditioned medium for 10 h (A, B) or 24 h (C) in the presence or absence of the DUB USP14 inhibitor IU1 (20 μM). The inhibitor IU1 was added 30 min prior and during OGD (preOGD), or immediately after (postOGD), and was present during the post-incubation period. Cell death was evaluated by nuclear morphology analysis (A,B), staining the nucleus with Hoechst 33342, and by the LDH activity present in the extracellular medium (C). The scale bar represents 50 µm. Full- length (280 kDa) and proteolytic fragments of α-spectrin (SBDPs) were analysed by western blot using an antibody recognizing both N- and C-terminal regions of the protein (D). The results are the average ± SEM of 4-5 independent experiments. Statistical analysis was performed by one- way ANOVA, followed by Bonferronís multiple comparison test, comparing selected pairs of columns (ns, p>0.5; *p˂ 0.05; **p˂ 0.01; ****p˂ 0.0001).

IU1 can have several pleiotropic effects through the regulation of known UPS targets (e.g mTOR), and more importantly, through the regulation of autophagy (Xu et al., 2015; Xu et al., 2016), all of them known to have an impact in neuronal survival in several ischemia/reperfusion models (Fletcher et al., 2013). To have a more direct evidence supporting the protective role of proteasome activation in cortical neurons subjected to OGD, we used two oleic amide derivatives that were recently shown to act as 20S proteasome activators (Halder et al., 2022). Compound 17 (renamed to B12) and compound 25 (renamed to D3) were used to ascertain their protective effects in cultured neurons subjected to OGD, and were both shown to boost the activity of the 20S proteasome by 0.5 and 3.5-fold, respectively (Halder et al., 2022). Cerebrocortical neurons were pre-incubated with the compounds (10 µM) for 2 h prior to the insult and then exposed to 1.5 h of OGD. Both compounds blocked OGD-induced cell death as determined by analysis of the nuclear morphology (Figure 7A,B; ns p>0.05). To ascertain the effect of the 20S proteasome activators on the cleavage of spectrin, we decided to use B12 since it showed the highest effect of the 20S proteasome (Halder et al., 2022). Similar to IU1, pre-incubation of the neurons with B12 also abrogated the effects of OGD on the activation of calpains, as determined by analysis of spectrin cleavage (Figure 7C; *p˂ 0.05; Figure 7. Supplementary information). Together, these results show restoring dysfunctional proteasomes is neuroprotective in the context of an in vitro model of ischemia and reperfusion.

**Figure 7.**
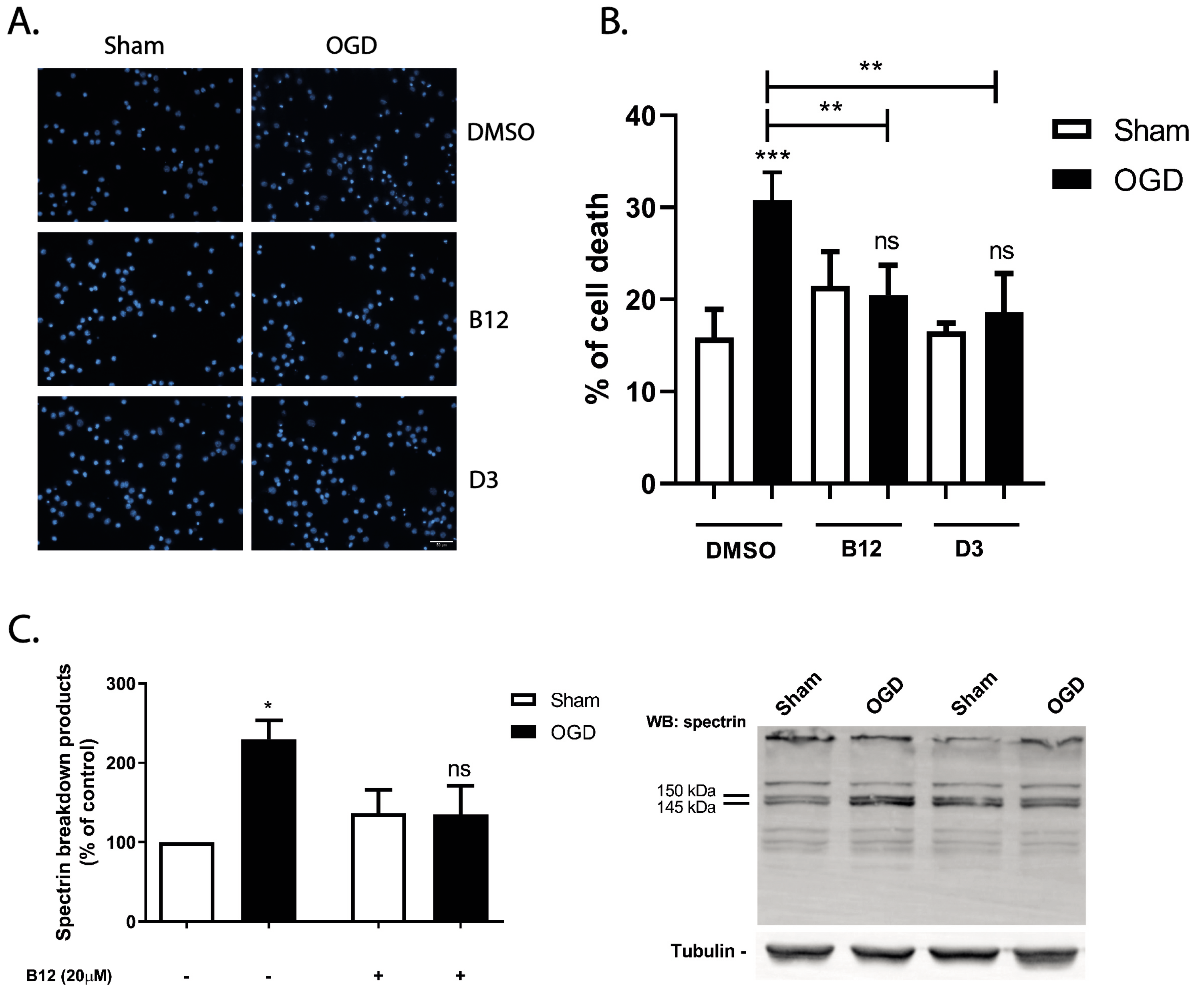
20S proteasome activators are protective against OGD-induced cell death. Cultured cerebrocortical neurons were subjected or not to OGD for 1.5 h, and further incubated in culture conditioned medium for 8 h in the presence or absence of the core particle activators B12 and D3 (10 μM). The activators were added 2 h prior and were present throughout the time of the experiments Cell death was evaluated by nuclear morphology analysis, staining the nucleus with Hoechst 33342 (A), and presented in the quantification graph as the percentage of cell death (B). The scale bar represents 30 µm. Full-length (280 kDa) and proteolytic fragments of α- spectrin (SBDPs) were analysed by western blot using an antibody recognizing both N- and C- terminal regions of the protein (C). The results are the average ± SEM of 4-5 independent experiments. Statistical analysis was performed by one-way ANOVA, followed by Bonferronís multiple comparison test, comparing selected pairs of columns (ns, p>0.5; *p˂ 0.05; **p˂ 0.01;***p˂ 0.001).

## Discussion

In this work we found that OGD-induced neuronal death is associated with calpain-dependent cleavage of proteasome subunits, together with the dissociation of the 20S catalytic core from the 19S regulatory particles. The impairment in the proteasome activity under these conditions is a key mediator of neuronal death as observed using two distinct experimental approaches to upregulate proteasome activity. Together, these results point to an important role for proteasome dysfunction in neuronal death in brain ischemia, which contrast with the protective effects of proteasome inhibitors observed in in vivo models of brain ischemia that have been attributed to the preservation of the blood-brain barrier (Doeppner et al., 2016; Doeppner et al., 2012; Williams et al., 2004; Williams et al., 2005; Zhang et al., 2006; Zhang et al., 2001).

Incubation of cultured cerebrocortical neurons under OGD for 1.5 h reduced the chymotrypsin-like activity of the proteasome by about 58%, when evaluated 4 h after the insult. In cultured hippocampal neurons OGD also induced time-dependent cell death by a mechanism dependent on the activity of NMDAR, resembling the mechanisms that contribute to neuronal death in brain ischemia (Caldeira et al., 2013). The decreased proteolytic activity of the proteasome after OGD correlates with its disassembly, as observed in cultured cortical neurons exposed to the same conditions. The decrease in the proteasome stability may result from the excessive stimulation of glutamate receptors, as suggested in experiments with hippocampal neurons subjected to excitotoxic stimulation (Caldeira et al., 2013). In accordance with the results obtained in in vitro experiments, transient global brain ischemia in rats was shown to impair the 26S proteasome activity, promote its disassembly and subsequent accumulation of polyubiquitinated proteins (Ge et al., 2007; Hochrainer, 2018). Together, these findings show that transient incubation of cultured neurons under conditions of OGD is a valuable model to investigate the molecular mechanisms involved in the disassembly of the proteasome in brain ischemia.

Using immunocytochemistry and antibodies specific for PMSA2 and Rpt6, commonly used as markers of the 20S catalytic particle and the 19S regulatory particle, respectively, we found that OGD has a differential effect on the dendritic distribution of the two proteasomal components. This is also in accordance with the results showing a disassembly of the proteasome in cultured cerebrocortical neurons subjected transiently to OGD. Previous work from our laboratory also showed that the nuclear and cytoplasmic population of proteasomes are differentially affected by excitotoxic glutamate stimulation, with the nuclear fraction being more affected under these conditions (Caldeira et al., 2013). Using a different model, it was shown that depriving yeast cells from nitrogen, but not glucose, induces proteasome disassembly followed by nuclear export and targeting to the vacuoles. However, the core and regulatory particles were targeted to the vacuoles in a differential manner, since only the former proteasomal component depended on the deubiquitinating enzyme Ubp3 (Waite et al., 2016). Despite the differences between neurons and yeast cells, it may be hypothesized that the clustering of Psma2 along dendrites at an early stage following OGD (2 h), in regions close to the synapse, may result from an increased trafficking of 20S proteasomes (without discarding putative effects on 26S proteasomes) to hotspots where there is an increased formation of reactive oxygen species. This may be relevant from the functional point of view since the 20S proteasomes are more effective in degrading oxidatively damaged proteins, another common hallmark of the ischemic brain (Daniele et al., 2021). The delayed increase in the Rpt6 puncta intensity after OGD in cortical neurons may suggest that regulatory mechanisms are induced to restoring normal and basal ubiquitin-dependent proteolysis after the 20S proteasome degrades the oxidatively damaged proteins. However, we can only speculate that the differential sorting of the 19S regulatory particle can serve as signaling complex by processing non-proteasome-targeting ubiquitin linkage (Sun et al., 2023).

The proteasome disassembly observed in in vivo models of brain ischemia as well as in cultured neurons subjected to OGD may allow the degradation of unfolded and oxidized proteins, which was shown to be carried out more effectively by 20S proteasomes, by a mechanism that does not require ubiquitination (Aiken et al., 2011; Shringarpure et al., 2003). In fact, we found that B12 and D13, two activators of the 20S proteasome, protected cultured cerebrocortical neurons subjected to OGD. Yet, whether these compounds activate already functional or latent proteasomes is still unknown and awaits further investigation (Zabel et al., 2010). Given the rather short post-incubation period used in this work, we cannot exclude the possibility that the disassembled proteasomes may reassemble at later time points, along with a recovery in the proteolytic activity. Together, the results suggest that proteasome disassembly may have a dual impact in cell survival. The concomitant increase in 20S proteasomes may allow to efficiently degrade oxidatively damaged proteins, in an unregulated and energy-independent manner, to cope with cellular stress (Sahu et al., 2021). However, an overall hypofunction of the proteasome may have deleterious consequences contributing to neuronal cell death.

Herein, we provide evidence showing calpain activation in cerebrocortical neurons subjected to OGD (1.5 h), when evaluated 4 h after the insult, as determined by an increase in the levels of spectrin-breakdown products of 150/145 kDa. The apparent absence of the 120 kDa SBDP under the same conditions rules out the possible contribution of caspase-3 to the cellular responses to the injury. The results are in accordance with other reports showing calpain activation in cultured hippocampal neurons subjected to OGD, when evaluated at different post-incubation periods after the insult (Curcio et al., 2015; Fernandes et al., 2014). We also showed that NMDAR act upstream of calpain activation in cerebrocortical neurons subjected to OGD, by using pharmacological tools to inhibit NMDAR and calpains. Under these conditions, no spectrin cleavage was observed, corroborating previous in vitro and in vivo findings (Curcio et al., 2015; Fernandes et al., 2014; Kawamura et al., 2005; Neumar et al., 2001; Roberts-Lewis et al., 1994). Numerous reports have identified calpain targets which are cleaved in brain ischemia thereby contributing to neuronal demise (Curcio et al., 2016). The experimental setup used in this work allowed the identification of two proteasome proteins that are cleaved by calpains, providing an insight about the molecular mechanism contributing to proteasome hypofunction in cortical neurons exposed transiently to OGD. In accordance with a previous report, we observed the calpain-mediated cleavage of the Rpn10 ubiquitin acceptor in cerebrocortical neurons subjected to OGD (Huang et al., 2013). Among the proteasome subunits tested, we also validated Rpt3 as a calpain substrate, using western blot experiments, along with Rpn10.

Although we have not investigated the functional impact of the calpain-mediated cleavage of proteasome subunits, there are several lines of evidence suggesting that mutations and knock- down of specific proteasome subunits affects its overall function. Thus, knockdown of the Rpt1, Rpn10, Rpn2 and Rpn12 proteasome subunits among others, reduced the viability of Schneider 2 (S2) cells, as well as the chymotrypsin-like activity (Wojcik and DeMartino, 2002). However, after glycerol-gradient centrifugation to isolate the different subpopulations of proteasomes, the activity of the 26S proteasome was only impaired when Rpn2, Rpt1 and Rpn12 were downregulated, whilst Rpn10 knockdown showed the opposite effect (Wojcik and DeMartino, 2002). Therefore, a reduction in Rpt1, Rpn2 and Rpn12 total protein levels has a profound effect on proteasome assembly. Additionally, the C-terminal region of the Rpt3 protein is essential for a proper assembly of the proteasome [(Kumar et al., 2010); see (Smith et al., 2007) for results showing the effect of point mutations and peptide entry], and turnover of polyubiquitinated proteins (Sokolova et al., 2015), whereas the Rpn6 protein was shown to interact with both Psma2 and Rpt6 protein, thus tethering the base of the 19S particle to the 20S (Pathare et al., 2012). In fact, its absence in yeast cells results in a severe impairment in the ubiquitin-proteasome pathway as shown by compromised assembled proteasomes and accumulation of polyubiquitin chains (Santamaria et al., 2003), whereas its overexpression increases proteasome assembly in embryonic stem cells (Vilchez et al., 2012). Rpn10-deficient mice showed embryonic lethality, and UIM domain deletion in this protein impaired the ubiquitin-dependent degradation of proteins in the mice liver (Hamazaki et al., 2007). Studies using a similar mice model with the UIM domain of Rpn10 genetically deleted showed an impaired interaction of polyubiquitinated substrates with the proteasome, as shown with purified proteasomes immunoblotted with an antibody against ubiquitin (Hamazaki et al., 2015). Genetic deletion of the Rpn10 ortholog in *Drosophila melanogaster* showed a dramatic increase in polyubiquitinated proteins, larval lethality along with an increase in 26S proteasomes [(Szlanka et al., 2003); see also (Wojcik and DeMartino, 2002)]. The available evidence, including the result showing Rpn10 cleavage by calpains (Huang et al., 2013), suggest that the stability/abundance of Rpn10 may not be the only factor responsible for controlling proteasome stability/assembly given the increase in proteasome activity when the protein is downregulated (Huang et al., 2013; Wojcik and DeMartino, 2002). However, the differential effects observed may be organism-and tissue-dependent and may also involve the degradation/interaction of other proteins.

Given the effects described above, the cleavage of Rpn10 alone is unlikely to account for the observed downregulation of proteasome activity. On the other hand, the cleavage of Rpt3 may contribute to the decrease in the stability of the proteasome after OGD (Kumar et al., 2010). Strikingly, one report showed that several calpain isoforms are in close proximity of the 26S proteasome, but whether there is a recruitment of these proteases to the vicinity of the 26S proteasome in the context of brain ischemia is unknown (Bartolome et al., 2024). It should also be considered that calpains cleave but do not degrade target proteins (Curcio et al., 2016). Therefore, the formation of truncated proteasome subunits may influence the proteasome structure and activity that is distinct from that obtained after deletion of the protein. Other factors that may also contribute to the regulation of the proteasome activity/stability after OGD include: (i) availability of ATP, which is important for the assembly of the 26S proteasome (Hendil et al., 2002; Liu et al., 2006); (ii) phosphorylation state of proteasome subunits (Bingol et al., 2010; Lokireddy et al., 2015).

The DUB USP14 acts as a brake in the degradation of proteasome substrates in vivo since its genetic deletion increased the overall protein turnover in yeast cells (Hanna et al., 2006), and similar results were obtained upon inhibition of the enzyme in cultured mammalian cells (Lee et al., 2010; Ponnappan et al., 2013). We found that preincubation of cortical neurons with the USP14 inhibitor IU1, as well as with the 20S proteasome activators B12 and D3, abrogates OGD-induced cell death. Interestingly, there was a decrease in calpain activity under the same conditions, suggesting that IU1 may act upstream of calpain activation, or it may reduce its activity through post-transcriptional regulation. In accordance with our observations, IU1 was reported to reduce brain infarct volume in a mouse model of transient focal cerebral ischemia at day 4 after the insult. In the latter study, elevation of the microRNA124 was found to downregulate USP14, and inhibition of this DUB mimicked the effect obtained under the former conditions (Doeppner et al., 2013). However, the possibility of an increased ubiquitin-dependent protein turnover was not considered in this study. More recently, two studies addressed the role of USP14 in controlling proteolysis in the cells, and showed an even more complex mechanism than initially proposed. USP14 is responsible for trimming both Lys48 and Lys63 polyubiquitin chains, which is in accordance with the consequent increase in the overall accumulation of polyubiquitin conjugates (Lee et al., 2010; Lee et al., 2016; Vaden et al., 2015; Xu et al., 2015). The neuroprotective effects of the USP14 inhibition may also be attributed to increased autophagy (Xu et al., 2016), being consistent with the effect of autophagy in neuroprotection in in vivo and in vitro ischemia/reperfusion models (Zhang et al., 2013). Because IU1 increases the turnover of proteins degraded in the UPS, identifying the putative candidates that are downregulated and contribute to the observed neuroprotective effects may provide an additional strategy for treating brain injury in ischemia/reperfusion models. Despite the multitude of mechanisms that may underlie the protective effects of IU1 in cerebrocortical neurons exposed to OGD, the increased cell survival also observed in the presence of the 20S proteasome activators clearly show a protective role of proteasome stimulation under the conditions used.

The results showing an inhibition of calpain activation by IU1 and by activators of the 20S proteasome in cortical neurons subjected to OGD, suggesting that the modulation of this proteolytic pathway may contribute to the protective effects of the DUB inhibitor. Since calpains cleave several proteasome subunits, as shown in this work, inhibition of these proteases in cells pretreated with IU1 or with the proteasome activators may enhance proteasome stability and protein degradation by the UPS, thereby stabilizing the proteostasis mechanisms. The protective effects of IU1 were only observed when the drug was present during OGD, suggesting that the maintenance of the proteostasis through activation of the UPS is relevant within a critical time period. However, the time required to achieve an effective concentration of IU1 inside the cells may prevent the effect of the DUB inhibitor when administered after OGD.

The impairment of the proteasome activity associated with neuronal death after OGD is in apparent contradiction with the reports pointing to a neuroprotective effect of proteasome inhibitors in several in vivo models of transient global/focal ischemia (Doeppner et al., 2016; Doeppner et al., 2012; Williams et al., 2004; Williams et al., 2003; Williams et al., 2005; Zhang et al., 2010; Zhang et al., 2006; Zhang et al., 2001). However, the latter results have been attributed mainly to the inhibition of the inflammatory responses resulting from the ischemic insult and to the stabilization of the blood-brain barrier. In fact, chemical proteasome inhibition is sufficient to increase neuronal cell death by inducing ER stress and activating the mitochondrial apoptotic pathway (Caldeira et al., 2013; Guo et al., 2016; Nishitoh et al., 2002). Therefore, therapeutic strategies to augment the degradative capacity of the UPS may also afford neuroprotection in several brain disorders.

In conclusion, we hereby propose a mechanism in which 30/26S proteasome disassembly and downregulation may be critical events downstream of calpain activation following transient brain ischemia leading to neuronal death. In addition, proteasome activation decreased OGD-induced calpain activation, showing a cross-talk between the two proteolytic systems. Proteasome activation in neurons is therefore a promising protective strategy in brain ischemia to complement the effects of proteasome inhibitors that act at the periphery to protect the integrity of the blood-brain barrier.

## Acknowledgements

This work was financed by the European Regional Development Fund (ERDF), through the Centro 2020 Regional Operational Programme and the COMPETE 2020 Operational Programme for Competitiveness and Internationalisation and Portuguese national funds via Fundação para a Ciência e a Tecnologia (FCT), under projects UIDB/04539/2020, UIDP/04539/2020 and LA/P/0058/2020.

